# Voluntary exercise during weight loss attenuates adipose CD8+ T cell exhaustion that persists after weight regain

**DOI:** 10.64898/2026.07.25.740730

**Authors:** Okechukwu Kingsley Oforka, Emily C. LaVoy, Matthew A. Cottam, Munira Kapadia, Emily Nguyen, Caroline Hegemann, Alyssa H. Hasty, Nathan C. Winn, Heather L. Caslin

**Author notes:** Corresponding authors: Heather Caslin, 3875 Holman St. Rm 104 Garrison, Houston, TX 77204-6015; Nathan Winn, 2215 Garland Avenue, 8465B MRBIV, Nashville, TN 37232.

## Abstract

**Background:** Weight gain and loss induce adipose CD8+ T cell exhaustion, which persists and may worsen glucose tolerance following weight regain. Because exercise can reduce T cell exhaustion in the blood, we hypothesized that exercise during weight loss would attenuate adipose CD8+ T cell exhaustion and glucose tolerance following weight regain.

**Methods:** Male C57Bl/6J mice were fed low-fat or high-fat diets over 8 to 9-week cycles to generate lean, obese, weight loss, or weight cycled groups. Additional weight loss and weight cycled groups were provided exercise wheels during the weight loss phase.

**Results:** As expected, weight loss increased total and exhausted CD8+ T cells by flow cytometry. Mice that ran the most during weight loss had the lowest proportion of exhausted CD8+ T cells. Notably, exercise reduced the proportion of exhausted CD8+ T cells even after the cessation of exercise and weight regain in all mice. However, exercise did not improve glucose tolerance or macrophage inflammation following weight regain. Moreover, exercise did not affect the induction of innate immune memory in adipose macrophages following weight loss.

**Conclusion:** The addition of exercise to a weight loss intervention remarkably reduced exhausted CD8+ T cells in the adipose tissue even after the cessation of exercise and weight regain. While exercise did not affect macrophage inflammation or glucose tolerance following weight regain, these results illuminate new questions about the persistence and mechanisms by which exercise reduces tissue CD8+T cell exhaustion and the direct role of macrophages in modulating glucose tolerance with weight cycling.

## 1. Introduction

Obesity increases one’s risk for many diseases, including cardiometabolic diseases, like diabetes^1,2^. These risks can be reduced with weight loss, but even when weight loss is initially successful, most people regain weight within a short period of time^3,4^. The widespread use of anti-obesity medications may further increase the prevalence or clinical importance of weight regain, as many individuals experience substantial weight regain after stopping treatment^5,6^. Cycles of weight gain, loss, and regain are referred to as weight cycling. Importantly, weight cycling worsens the risk of many cardiometabolic diseases even more than obesity itself^7,8^. While the mechanisms linking obesity to worsened disease outcomes have been broadly studied, the mechanisms underpinning the link between weight cycling and disease have received less attention.

Metabolic abnormalities associated with weight cycling may arise from aberrant immune responses. With obesity, adipose tissue T cells increase in number, drive local macrophage recruitment and inflammation, and worsen glucose tolerance^9^. Adipose expansion also increases CD8+ T cell exhaustion^10^, characterized by impaired effector function and sustained expression of inhibitory receptors^11^. While weight loss normalizes systemic metabolism, many immune populations persist or even increase following weight loss, including many T cell populations^12^. Specifically, diet-induced weight loss and weight regain further increase the percentage of adipose CD8+ T cells, including memory CD8+ T cells with an exhausted phenotype^12,13^. Recent data suggests that blocking CD8+ T cell activation and memory development through CD70 improves the glucose impairments specifically related to weight regain^14^. Thus, CD8+ T cell exhaustion may be a potential target to improve glucose tolerance with weight cycling.

Exercise represents a potential intervention for weight cycling capable of targeting both metabolic and immune pathways^15^. Regular exercise seemingly reduces exhausted CD8+ T cells in the blood after acute exercise and chronic exercise training^16,17^. In addition, exercise training improves glucose tolerance and reduces the risk of developing type 2 diabetes^18^. Therefore, we aimed to A) determine the effect of exercise on adipose CD8+ T cell exhaustion following weight gain and loss and B) the impact of exercise on adipose CD8+ T cell exhaustion and glucose intolerance amplified by weight cycling. We hypothesized that the addition of voluntary wheel running to diet-induced weight loss would reduce CD8+ T cell exhaustion and glucose intolerance worsened by weight regain.

## 2. Methods

### 2.1. Ethics Statement

All studies were approved and completed in compliance with the University of Houston Institutional Animal Care and Use Committee (IACUC; PROTO202300026) and the Vanderbilt University IACUC (V1800143). The University of Houston and Vanderbilt University are accredited by the Association for Assessment and Accreditation of Laboratory Animal Care International.

### 2.2. Murine models of weight loss and weight cycling

For all experiments, C57Bl/6 male mice were put on 8-9-week cycles of low-fat (10% fat, Research Diets, #D12450B/J) or high-fat diets (60% fat, Research Diets, #D12492) around 7–8 weeks of age (see experiment schematics: Fig 1A, Fig 2A, Fig 3A). In all voluntary wheel running groups, mice were permitted to exercise with access to freely moving Prevue Hendryx E-Z Roller Exercise Wheels- 4.5’ as detailed in experimental schematics. Wheel run distance was measured approximately each day using a Velo 7CATEYE Wired Bike Computer or Artyea KINGMAS LCD Bicycle Bike Computer Odometer Speedometer Sd-548b. To control for enrichment, non-exercise mice had access to stationary wheels. For all mice, food was provided ad libitum, and body weight was recorded weekly.

**Figure 1:**
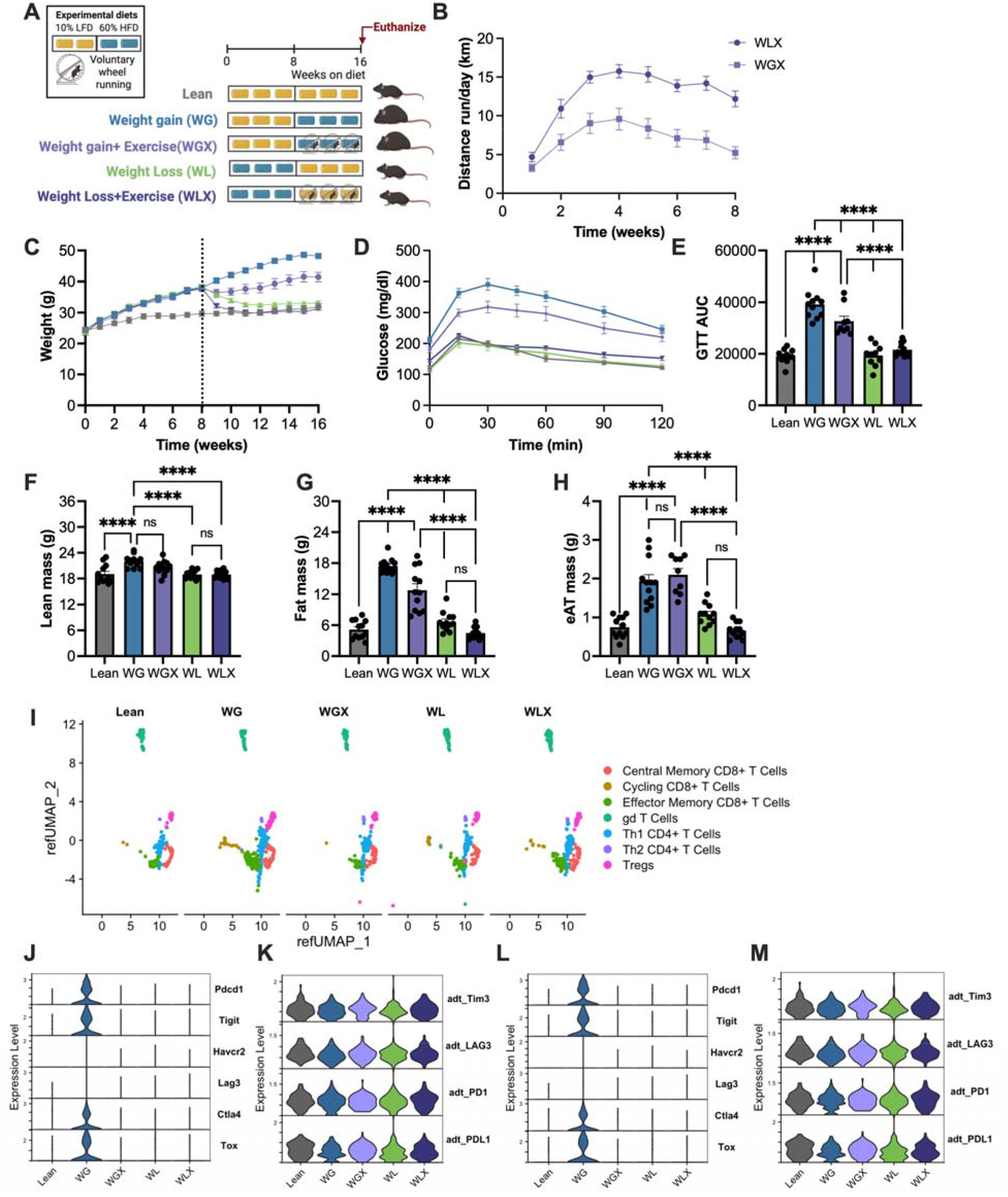
Exercise seemingly reduces exhaustion proteins on CD8+ T cells with weight gain and weight loss. A) Schematic: Male C57Bl/6 mice were placed on cycles of low-fat and/or high-fat diets for 16 weeks to elicit lean, weight gain (WG), and weight loss (WL) mice. Two additional groups underwent voluntary wheel running (WGX and WLX). Figure schematic was created using Biorender.com. B) Distance run per day was recorded for the WGX and WLX mice during the exercise period (second 8 weeks). C) Body weight measured weekly over 16 weeks. D) Blood glucose over time and E) area under the curve (AUC) for a glucose tolerance test (GTT; 1.5 g/ kg lean mass). F) Lean mass and G) fat mass were measured by EchoMRI in week 17. H) Epididymal adipose tissue (eAT) mass was measured following euthanasia. Means ± SEM, ****p<0.001 by one-way ANOVA and post-hoc analysis. T cell populations were subset from the single cell RNA-sequencing data and I) a Uniform Manifold Approximation and Projection (UMAP) of these populations shows the distribution by group. Expression of exhaustion related genes and proteins is shown for J&K) all CD8+ T cells and L&M) CD8+ memory T cells, respectively.

**Figure 2:**
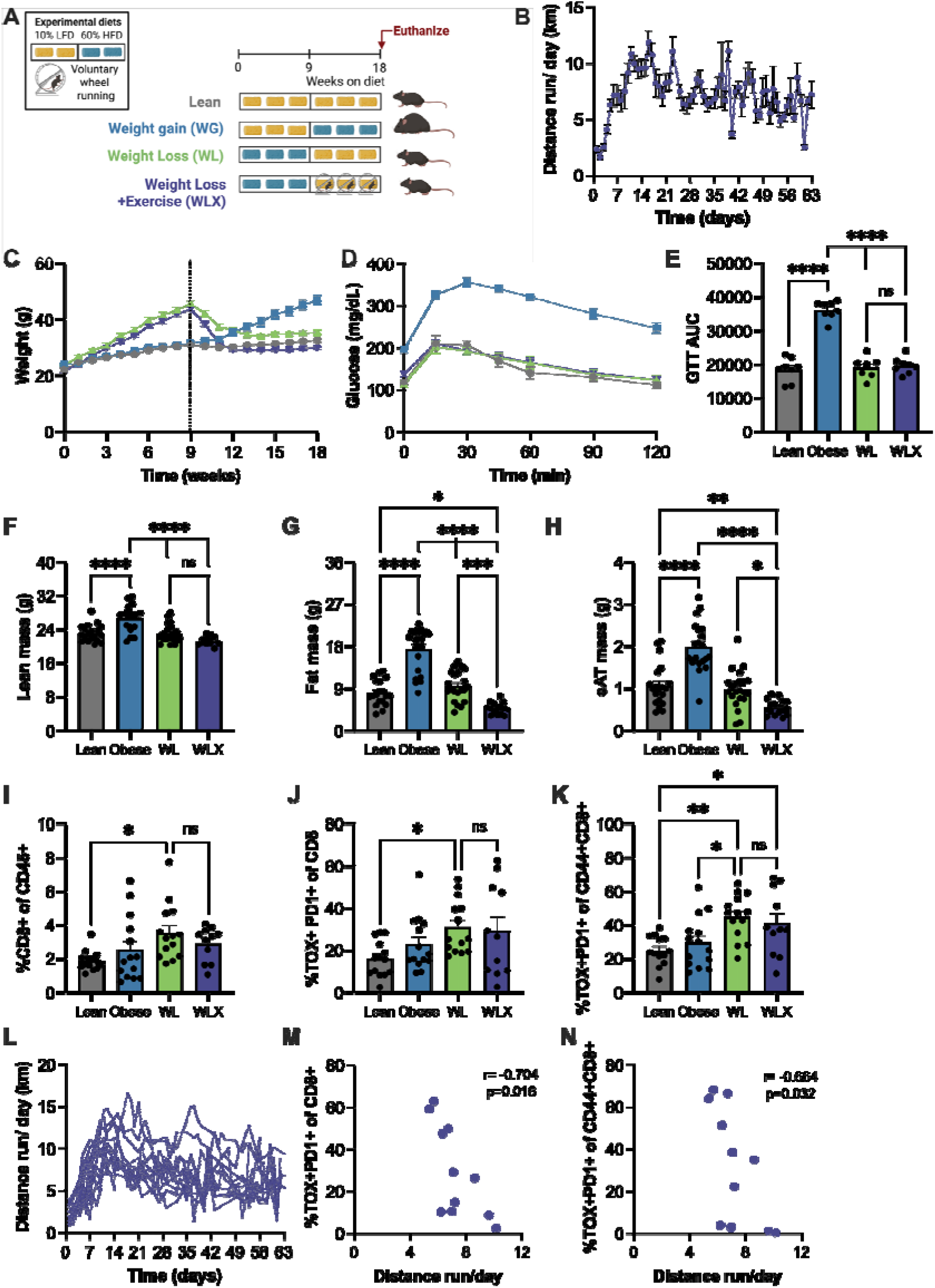
Exercise during weight loss correlates with reduced CD8+ T cell exhaustion in the adipose tissue. A) Schematic: Male C57Bl/6 mice were placed on cycles of low-fat and/or high-fat diets for 18 weeks to elicit lean, obese, and weight loss (WL) groups. One additional weight loss group underwent voluntary wheel running during weight loss (WLX). Figure schematic was created using Biorender.com. B) Distance run per day was recorded for the WLX mice during the exercise period (second 9 weeks). C) Body weight measured weekly over 18 weeks. D) Blood glucose over time and E) area under the curve (AUC) for a glucose tolerance test (GTT; 1.5 g/ kg lean mass). F) Lean mass and G) fat mass were measured by EchoMRI in week 17. H) Epididymal adipose tissue (eAT) mass was measured following euthanasia. I) % CD8+ T cells (CD45+TCRb+CD8+), J) %TOX+ PD-1+CD8+ T cells, K) %TOX+ PD-1+ of Memory CD8+ T cells (CD44+CD8+ T cells) were measured by flow cytometry. L) Distance run per day was graphed individually and M) %TOX+ PD-1+CD8+ T cells and N) %TOX+ PD-1+ of Memory CD8+ T cells were correlated with distance run per day for each mouse. Means ± SEM, *p<0.5, ***p<0.01, ****p<0.001 by one-way ANOVA and post-hoc analysis (or correlation matrix for L&M).

**Figure 3:**
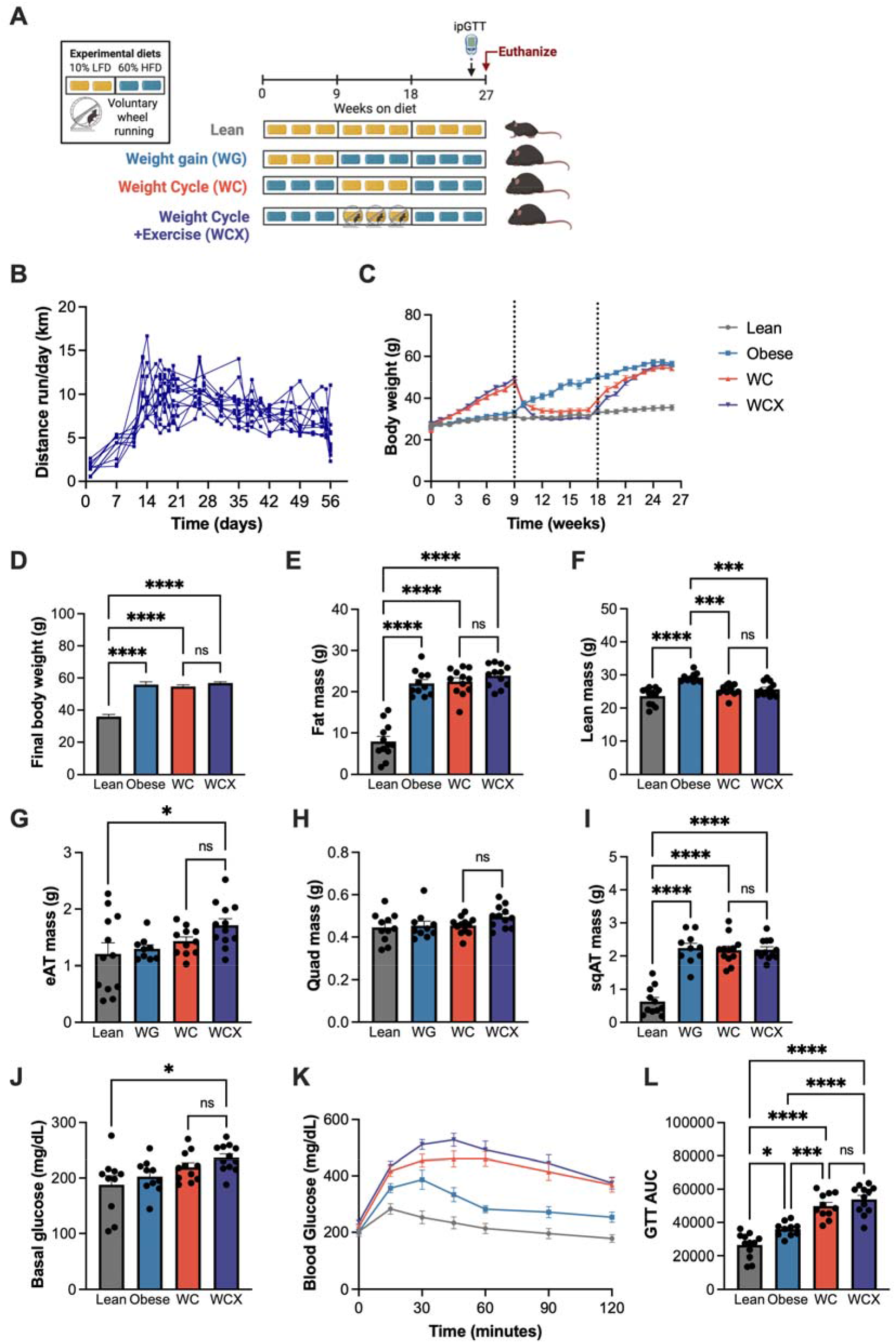
Exercise during weight loss does not improve glucose tolerance following weight regain. A) Schematic: Male C57Bl/6 mice were placed on cycles of low-fat and/or high-fat diets for 27 weeks to elicit lean, obese, and weight cycled (WC) groups. One additional weight cycling group underwent voluntary wheel running during weight loss (WCX). Figure schematic was created using Biorender.com. B) Distance run per day was recorded for the WCX mice during the exercise period (second 9 weeks). C) Body weight measured weekly over 27 weeks. D) Final body weight measured at week 27th. E) Lean mass and F) fat mass measured by EchoMRI in week 26. G) Epididymal adipose tissue (eAT) mass, H) quadriceps mass, I) subcutaneous adipose tissue (sqAT) mass were measured following euthanasia. J) Basal glucose, K) blood glucose over time, and L) blood glucose area under the curve (AUC) for a glucose tolerance test (GTT; 1g/ kg total mass) in week 27. Means ± SEM, *p<0.5, ***p<0.01, ****p<0.001 by one-way ANOVA and post-hoc analysis.

### 2.3. Body composition and glucose tolerance testing

To determine the effect of weight loss, weight regain, and exercise on body composition, lean and fat mass were measured one week prior to the end of each study by EchoMRI (University of Houston) or nuclear magnetic resonance (Bruker Minispec; Vanderbilt University).

To determine the effect of weight cycling and exercise on glucose tolerance, an intraperitoneal glucose tolerance test was administered one week prior to the end of the study. A 20% dextrose solution was prepared, and the required volume was calculated to deliver 1.0 g dextrose/kg of total body mass or 1.5g dextrose/ kg lean mass (specified in figure legends). Following baseline sampling, mice received an intraperitoneal injection of the calculated dextrose dose. Blood glucose was measured at 15, 30, 60, 90, and 120 min post-injection via glucometer (CVS Advanced glucometer or Contour next ONE).

At the conclusion of each study, mice were euthanized and epididymal adipose tissue was weighed. In the weight cycling study, subcutaneous adipose tissue and quadriceps mass were also measured.

### 2.4. Tissue immune cell isolation

To determine the impact of weight loss, weight cycling, and exercise on immune parameters, the stromal vascular fraction (SVF) was isolated from epididymal adipose tissue using established techniques^19^. Following euthanasia, phosphate buffered saline (PBS) was perfused through the vasculature. Adipose tissue was weighed and then minced with scissors, enzymatically digested with type IV collagenase (Worthington Biochemical, #LS004176), filtered through 100 µm filters, and centrifuged to remove adipocytes. Samples were then incubated with ACK lysis buffer (Quality Biological, #118-156-101) for red blood cell removal. SVF was then used for downstream single cell RNA-sequencing and flow cytometry analysis.

### 2.5. Single cell RNA-sequencing

To comprehensively assess immune populations in the SVF, we used single cell RNA-sequencing in 4 mice/ group with cellular indexing of transcriptomes and epitopes by sequencing (CITE-seq) to support protein-level changes like in our previous publications^12,14^. First, Fc Block (BD Biosciences, #553141553141) was added to prevent non-specific binding. Cells from each mouse were then labeled with unique hashtag antibodies (TotalSeq-C, Biolegend; see Supplemental Table 1) and anti-CD45 microbeads (Miltenyi Biotec, #130-110-608) per manufacturer instructions, and biological replicates were pooled and sorted on the manual Miltenyi Biotec OctoMACS Separator for immune isolation. The resulting CD45+ cells were then labeled for CITE-seq using TotalSeq-C antibodies for cell-surface markers to identify major cell types and T cell phenotypes (see Supplemental Table 1).

Sample preparation was conducted using the 5′ assay for the 10X Chromium platform (10X Genomics), targeting 20,000 cells per diet group (∼5,000 cells/ sample). In all, 50,000 reads per cell were targeted for PE-150 sequencing on the Illumina NovaSeq6000.

### 2.6. Sequencing data processing

FastQ files obtained from sequencing were processed using CellRanger (version V4; 10X Genomics) with feature barcoding. CellRanger outputs were further processed using the R package SoupX V1.5.283 to remove ambient contaminating RNA^20^. The Seurat V4 R package was used for quality control, data set integration, clustering, cell type annotation, differential expression, and visualization^21^. Strict quality control parameters were utilized post sequencing to ensure that only viable cells which could be traced back to each individual mouse were used for further analysis. For quality control of sequenced cells, only cells that had at least 200 gene features, at least 500 total measured RNA sequences, and <5% mitochondrial RNA content were retained. Furthermore, only cells designated as singlets after hashtag demultiplexing (i.e. cells containing at least one hashtag, but not more than one) were used for visualization and analysis. DoubletFinder V3 was used to detect heterotypic doublets and further confirm singlets^22^. Harmony V0.1 was used for batch-aware integration for datasets sequenced at different times^23^.

Cell type annotation was performed using reference mapping with the *MapQuery* function in Seurat V4 to our previously published murine adipose immune cell atlas^12^ and confirmed using canonical cell type markers. Cell clustering can be visualized in Supplemental Fig 1A and cluster identities can be confirmed in Supplemental Fig 1B. All downstream gene analyses were performed using the normalized RNA assay.

### 2.7. Data availability

The single cell RNA-sequencing data from this study have been submitted to the National Center for Biotechnology Information Gene Expression Omnibus (GEO) and will be made available upon publication. The GEO accession number is GSE341256.

### 2.8. Flow cytometry

Flow cytometry was used to directly characterize T cell and macrophage changes in the SVF in both weight loss and weight cycling studies. Zombie violet stain (Biolegend, #423114) was added for 15 min to assess viability. Following a 10-minute incubation with TruStain FcX™ (anti-mouse CD16/32; Biolegend, #101320), cells were stained for extracellular protein surface markers with antibody panels as in Supplemental Table 2 for 45 minutes at 4°C. For intracellular transcription factors, cytokines, and histone modifications, samples were then fixed and permeabilized with the Foxp3/Transcription Factor Staining Buffer Set (Invitrogen/eBioscience, #00552300) according to the manufacturer’s protocol and intracellular target antibody staining occurred overnight at 4°C. Cells were washed once and data were collected on a Miltenyi MACSQuant 10 flow cytometer and analyzed using FlowJo software (Version 10). See Supplemental Figs 2&3 for gating strategy.

### 2.9. Statistical analyses

Data were analyzed and graphed using GraphPad Prism (V10). Outliers were removed, and data was tested for assumptions of normality (Shapiro Wilk test and visualization of the Q-Q plot). Data with a normal distribution were analyzed using a one-way ANOVA to examine group differences and Tukey’s multiple comparisons test for post-hoc analyses where appropriate. Data with a non-normal distribution were analyzed using a Kruskal-Wallis test with Dunn’s multiple comparison post-hoc tests as appropriate. All statistical analyses were performed with statistical significance set at p < 0.05.

## 3. Results

### 3.1. Exercise reduces CD8+ T cell exhaustion following weight gain and weight loss

Exercise has been previously shown to reduce the proportion of exhausted T cells in the blood^16,17^. We have also previously shown that weight loss further induces CD8+ T cell exhaustion beyond obesity^12^. Thus, we aimed to determine if exercise can reduce CD8+ T cell exhaustion in the adipose tissue following weight loss. Mice were put high-fat diets for 8 weeks and then maintained on high-fat diet with or without access to freely moving wheels (WG and WGX) or switched to low-fat diet with or without access to freely moving wheels (WL and WLX; Fig 1A). A final group was maintained on low-fat diet throughout (lean). Distance run is shown in Fig 1B. At the end of the study, WG mice had the highest body weight as expected, and voluntary wheel running reduced body weight in the WGX vs WG groups (Fig 1C). The lean, WL, and WLX groups were all the same weight at 18 weeks. WG mice had worse glucose tolerance than lean mice (Fig 1D&E). WGX significantly reduced glucose impairments compared to WG mice. WL and WLX restored glucose tolerance to that of lean controls. Lean mass was highest in the WG group, though not significantly different from WGX (Fig 1F). Total fat mass and epididymal adipose tissue mass were both highest in WG mice and reduced with WL (Fig 1G&H), and WGX reduced total fat mass compared to WG mice.

T cell populations were assessed by single cell RNA-sequencing for each group (Fig 1I). Interestingly, while WG mice showed the highest expression of the exhaustion related genes *Pdcd1, Tigit, Ctla4*, and *Tox* in all CD8+ T cells (Fig 1J) and memory CD8+ T cells (Fig 1L), CD8+ and CD8+ memory cells in the weight loss group showed the highest expression of TIM3, LAG3, PD-1, and PD-L1 by protein expression (Fig 1K&M, respectively). By protein expression, TIM3, LAG3, PD-1, and PD-L1 were all reduced in the WLX compared to the WL group. Together, these data suggest exercise during weight gain and weight loss reduces CD8+ T cell exhaustion in the adipose tissue.

### 3.2. Reductions in CD8+ T cell exhaustion correlate with exercise volume during weight loss

We were particularly interested in understanding how the inclusion of exercise during a weight loss intervention could affect immune populations and glucose tolerance following weight regain. Thus, we conducted two replicate experiments in which mice were put on low-fat and high-fat diets for intervals of 9-weeks to correspond with the 18-week timepoint of our well-established model of weight cycling^12–14,24^. Additionally, for this experiment, adipose T cells were analyzed by flow cytometry, which allowed for a more direct quantification of CD8+ T cell number and proportion and the expression of surface proteins at a single cell level. Thus, we generated weight loss (WL) mice with lean and obese controls. The obese and WL mice matched for total time on high-fat diet (Fig 2A). A fourth group underwent diet-induced weight loss and was provided access to freely moving running wheels (WLX). Distance run is shown in Fig 2B. At 18 weeks, lean, WL, and WLX mice did not significantly differ in body weight (Fig 2C).

Glucose tolerance was measured in one cohort of mice, and as previously published^12^, WG worsened glucose tolerance compared to lean mice, and this was rescued with WL (Fig 2D&E). There was no significant difference in glucose tolerance between lean, WL, and WLX mice. Moreover, like the experiment from Fig 1, lean mass was highest in the obese group (Fig 2F). Both total fat mass and epididymal adipose tissue mass were highest in WG mice and there was a significant reduction in both adipose measures in the WLX group compared to WL (Fig 2G&H).

To assess the adipose T cell compartment, we used flow cytometry for a better understanding of total cell numbers, proportions, and phenotypes. The number of CD8+ and memory (CD44+) CD8+ T cells were increased in the WL group (Supplemental 4C&D), but there was no significant difference in the WL and WLX groups. Moreover, the % CD8+ T cells, exhausted CD8+ T cells (PD-1+TOX+CD8+), and exhausted memory CD8+ T cells (PD-1+TOX+CD44+CD8+) also increased in the WL group but were not different between the WL and WLX groups (Fig 2I-K). This was seemingly different from the sequencing data; however, when further analyzed, it was noted that running distance/ day was quite variable in these cohorts (Fig 2L) and also lower than in Fig 1. Interestingly, % exhausted CD8+ and exhausted memory CD8+ T cells were significantly correlated with distance run/day (Fig 2M&N). There was no correlation observed between either T cell population and body weight or fat mass (data not shown). Thus, the reduction in adipose CD8+ T cell exhaustion with exercise is seemingly dependent on exercise volume.

### 3.3. Exercise during weight loss does not improve glucose tolerance following weight regain

Because exercise affected CD8+ T cell exhaustion after weight loss above, we wanted to determine if these effects would persist and affect metabolism after weight regain. We generated lean, obese, and weight cycled (WC) mice over 27 weeks as previously published (Fig 3A;^12–14,24^. One additional WC group engaged in voluntary wheel running during the nine-week weight loss phase (WCX), after which running was discontinued. Distance run is shown in Fig 3B.

At 27 weeks, body weight was increased in obese, WC, and WCX mice compared with lean controls (Fig 3C-D). Importantly, the obese, WC, and WCX mice were on high-fat diets for the same length of time and had similar body weight and fat mass at the end of the study (Fig 3D-E), allowing for interpretation of metabolic and immunological data by intervention and not solely due to adiposity. Lean mass was only elevated in the obese group (Fig 3F), and epididymal adipose tissue weight was only elevated in the WCX group (Fig 3G). There was no difference in weight of the quadriceps muscle (Fig 3H), and like total fat mass, subcutaneous adipose tissue weight was elevated in the obese, WC, and WCX groups (Fig 3I).

To assess glucose metabolism, we measured basal glucose levels and glucose tolerance. Basal (fasting) glucose was elevated in WCX mice compared to lean mice (Fig 3I). As previously published, obese mice showed impaired glucose tolerance compared with lean mice, and glucose tolerance was further impaired in WC mice (Fig 3K&L). Surprisingly, WCX did not show improved glucose tolerance compared to the WC group. Thus, the inclusion of exercise during diet-induced weight loss is not enough to protect against glucose intolerance following weight regain and the cessation of exercise.

### 3.4. Exercise during weight loss attenuates CD8⁺ memory T cell exhaustion following weight regain

We next tested if exercise during weight loss had a lasting impact on CD8+ T cell exhaustion following weight regain. Flow cytometric data from the epididymal adipose tissue showed that obese, WC, and WCX mice had a significant increase in the number of immune cells (CD45+) per gram of adipose tissue compared to the lean group as expected; however, WC was the only group to show a significant increase in the proportion of immune cells (Fig 4A). The number of adipose tissue T cells (TCRb+CD45+) also increased in the WC group, but the percent and number of total T cells was lower in the WCX vs. WC group (Fig 4B). Obesity and WC significantly increased the number of CD8+ and memory (CD44+) CD8+ T cells (Fig 4C&D), and prior exercise significantly attenuated the number and proportion of CD8⁺ T cells in the WCX vs. WC group (Fig 4C). Prior exercise also reduced the number of CD8+ memory T cells in the WCX vs. WC groups (Fig 4D), but there was no effect on cell proportion, likely due to sample variability. Exhausted (TOX+PD-1+) CD8+ T cells and exhausted memory CD8+ T cells were increased in obesity and WC (Fig 4E), as previously observed^12^. Interestingly, WCX mice had significantly less exhausted cells in both total and memory CD8+ T cell populations, despite having no impact on glucose tolerance. To further phenotype the CD8+ T cells, we also measured basal cytokine levels. While the percentage of IFN-γ+ CD8+ T cells did not differ among groups (Fig 4G), granzyme B expression decreased significantly in the obese and WCX groups (Fig 4H). WC and WCX groups did not differ in granzyme B expression. Thus, while exercise during weight loss did not prevent glucose intolerance with weight cycling, the reduction in CD8+ T cell exhaustion was sustained following the cessation of exercise and weight regain.

**Figure 4:**
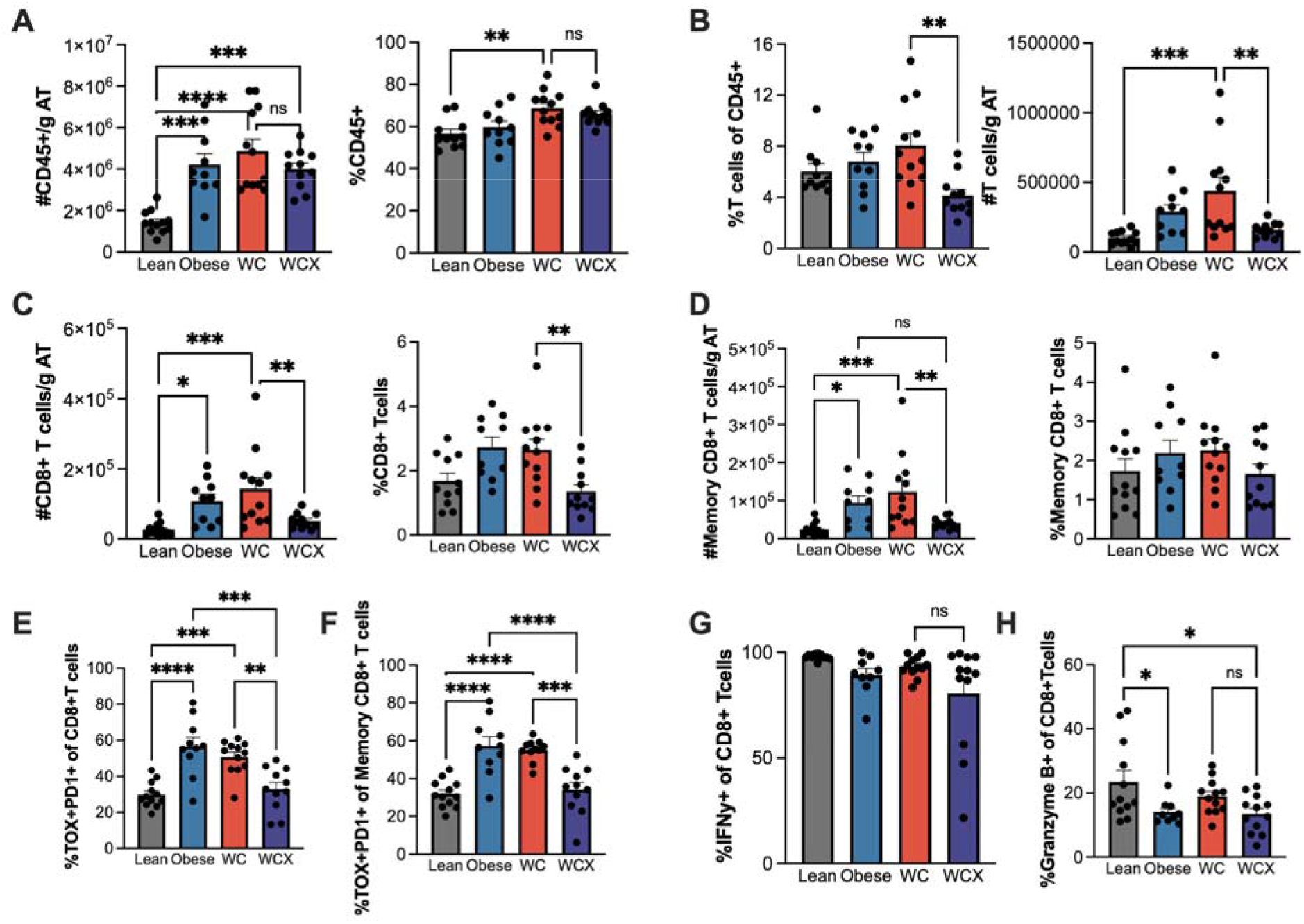
Exercise during weight loss attenuates CD8+ memory T-cell exhaustion with weight cycling. # of cells/ gram of epididymal adipose tissue (AT) and % for A) CD45+ immune cells, B) T cells (CD45+TCRb+), C) CD8+ T cells (CD45+TCRb+CD8+), and D) CD8+ Memory T cells (CD45+ TCRβ+CD8+CD44+) for lean, obese, weight cycled (WC) and weight cycle+ exercise (WCX) mice were quantified by flow cytometry. E) %TOX+ PD-1+ of CD8+ T cells F) %TOX+ PD-1+ of Memory CD8+ T cells, and G) %IFNγ+ CD8+ T cells, H) %Granzyme B+ of CD8+ T cells were also quantified. Means ± SEM, *p<0.5, ***p<0.01, ****p<0.001 by one-way ANOVA and post-hoc analysis.

### 3.5. Exercise during weight loss does not reduce macrophage inflammation following weight regain

Because exercise did not reduce glucose intolerance despite the reduction in exhausted CD8+ T cells, we aimed to determine if another cell population could better explain the glucose tolerance data. Macrophages in adipose tissue also play a central role in obesity-associated metabolic dysfunction, where they accumulate and secrete pro-inflammatory cytokines that contribute to insulin resistance^25^. We and others have shown that macrophage inflammation persists in weight loss and weight regain^24,26,27^. Thus, we also measured the impact of exercise on macrophage function in weight cycling. The number of myeloid cells (CD11b+CD45+) and macrophages (CD11b+F4/80+CD45+) per gram of epididymal adipose tissue were significantly increased in obese, WC, and WCX mice compared with the lean group (Fig 5A&B), yet the percent of each population was highest in the WCX group. To assess macrophage inflammatory status, basal expression of the proinflammatory cytokines TNF and IL-6 were quantified within the macrophage population. There was a significant increase in TNF+ and IL-6+ macrophages in obese mice compared to lean and a further increase in TNF+ and IL-6+ macrophages in WC and WCX mice (Fig 5C&D). WC and WCX did not differ in the proportion of TNF and IL-6-expressing macrophages. The relative amount of TNF and IL-6 by median fluorescence intensity (MFI) was also elevated in the obese and WC groups and remained high in WCX mice (Fig 5C&D). Together, these findings demonstrate that while prior exercise induces lasting reductions in CD8+ T cell exhaustion, macrophage inflammation persists in weight cycling and corresponds with sustained glucose impairments.

**Figure 5:**
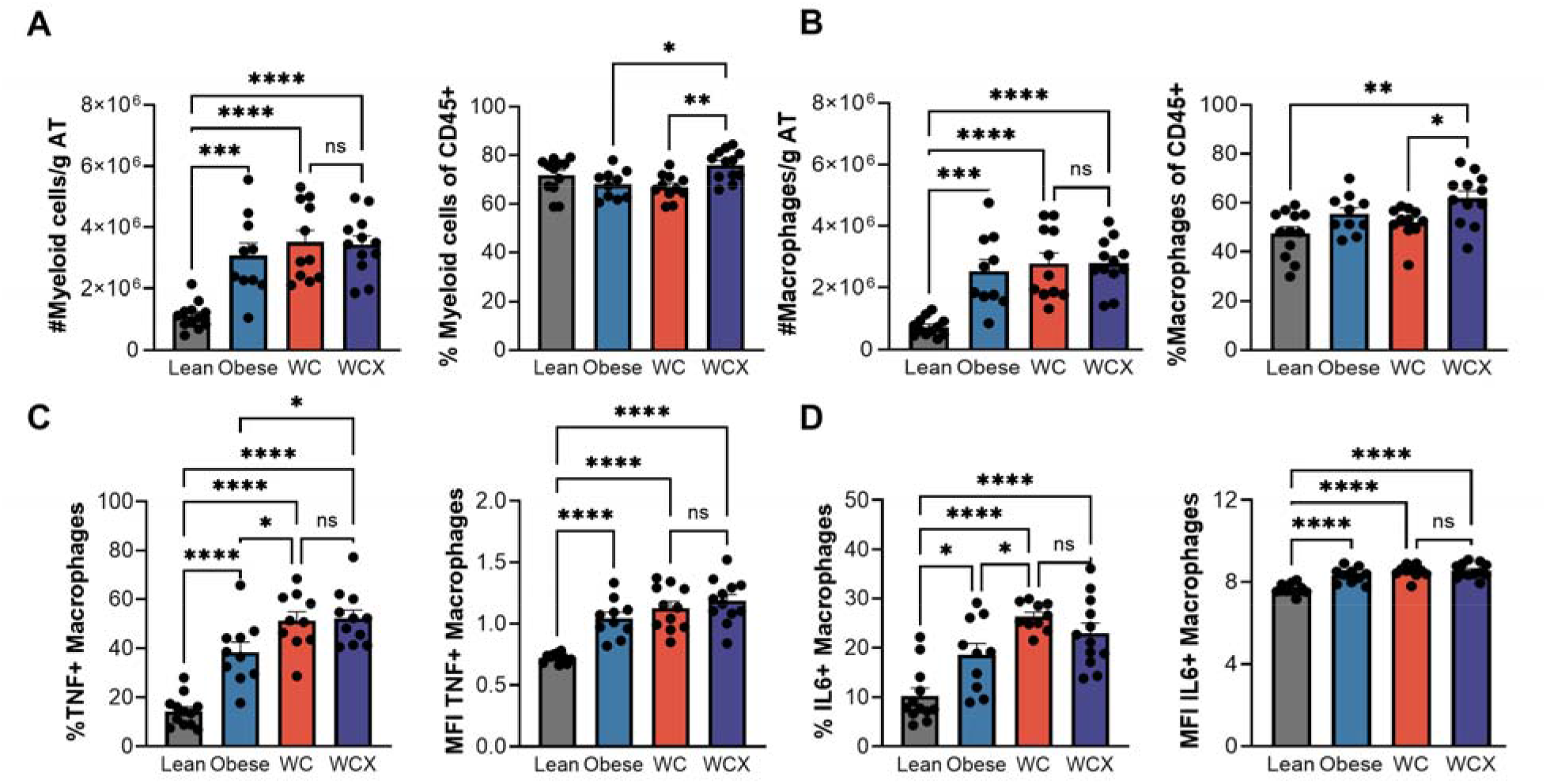
Exercise during weight loss does not reduce adipose macrophage inflammation with weight cycling. # cells/ gram of epididymal adipose tissue (AT) and % for A) myeloid cells (CD45+CD11b+) and B) macrophages (CD45+CD11b+F4/80+) for lean, obese, weight cycled (WC) and weight cycle+ exercise (WCX) mice were quantified by flow cytometry. % and median fluorescent intensity (MFI) of positive cells were quantified for C) TNF and D) IL-6 within the macrophage population. Means ± SEM, *p<0.5, **p<0.1, ***p<0.01, ****p<0.001 by one-way ANOVA and post-hoc analysis.

### 3.6. The addition of exercise to diet-induced weight loss does not prevent the induction of innate immune memory in adipose macrophages

Previous work from our group has shown that weight loss specifically induces innate immune memory in adipose tissue macrophages^24^. Innate immune memory is defined by the metabolic and epigenetic adaptations that enhance responsiveness to a subsequent stimulus^28^. To directly examine how exercise impacts innate immune memory in adipose tissue macrophages, we first returned to the macrophage data within our single cell RNA-sequencing data (see schematic from Fig 1A and Supplemental Fig 5A). At the gene expression level, WG increased the expression of the inflammatory gene *Tnf* and the metabolic genes *Hk3, Ldha, Sdhb,* which did not return to baseline with WL or exercise (Supplemental Fig 5B&C). These data suggest persistent changes in both inflammation and metabolism. Interestingly, *Acly* was elevated in WGX and WLX. As expected, by gene expression WG increased the proportion of many lipid handling genes, however, *Trem2, Fabp4, Plin2, Lipa, and Cd9* remained elevated specifically in the WLX group (Supplemental Fig 5D). Last, at the protein level, WLX had the highest expression of MHCII (Supplemental Fig 5E), together suggesting a memory of obesity, or even an increased inflammatory profile in the WL and WLX macrophages compared to macrophages from obese mice.

We also assessed macrophages by flow cytometry following two 9-week cycles of low-fat and/or high-fat in lean, obese, weight loss (WL), and weight loss+ exercise (WLX) groups (see schematic in Fig 2A). Immune cells were increased in number in the obese, WL, and WLX mice relative to lean, but only increased by % in the obese group (Fig 6A&B). The number of macrophages were elevated compared to lean in the obese, WL, and WLX groups, while again the % macrophages was elevated in the obese group only (Fig 6C). To understand how exercise may affect epigenetic modifications involved in innate immune memory, we assessed the % of macrophages which had high expression of H3K4me3 and H3K27ac. The % of H3K4me3 high macrophages was elevated in WL mice as previously published [^24^ and Fig 6D)], and remained elevated in WLX mice. Additionally, WL increased the % H3K27ac+ macrophages compared to lean and WG mice as previously published [^24^ and Fig 6E)]. However, the % H3K27ac+ macrophages did not differ between WL and WLX mice. Finally, we assessed macrophage cytokine production following *ex vivo* activation with LPS. Percent IL-6+ macrophages in the WL and WLX groups trended towards higher basal IL-6 production, but this was not significant (Fig 6F). Following LPS activation, the % TNF+ and IL-6+ macrophages were elevated in both the WL and WLX groups, with no difference between WL and WLX groups (Fig 6F&G). Together, this data suggests that the addition of exercise to a diet-induced weight loss intervention does not prevent the induction of innate immune memory in adipose macrophages.

**Figure 6:**
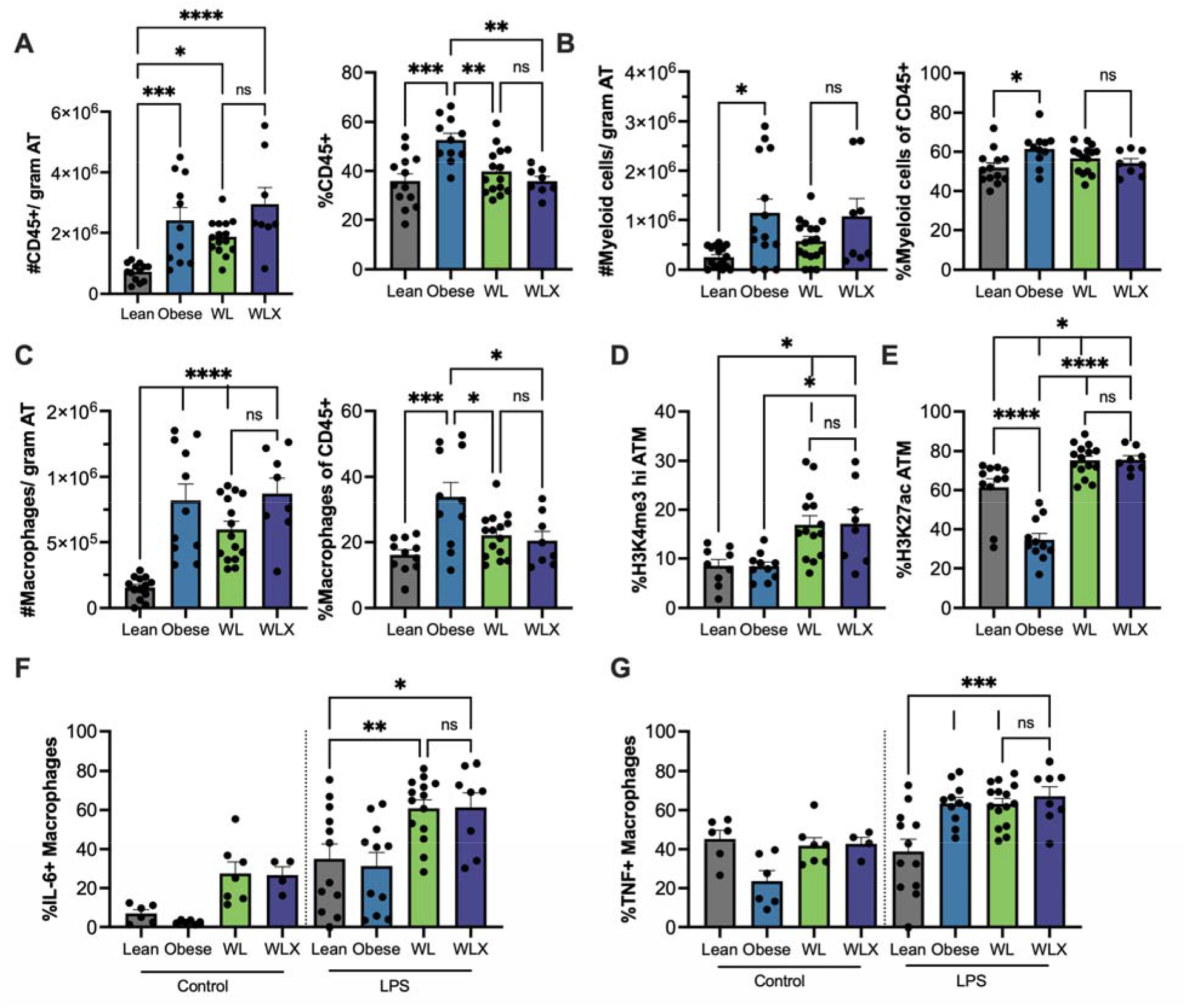
Exercise does not lessen adipose macrophage memory with weight loss. Using 18 week lean, obese, weight loss (WL), and weight loss + exercise (WLX) mice from Figure 2, # cells/ gram of epididymal adipose tissue (AT) and % for A) total immune cells (CD45+), myeloid cells (CD45+CD11b+) and C) macrophages (CD45+CD11b+F4/80+) were quantified by flow cytometry. D) %H3K4me3 high and E) H3K27ac epigenetic modifications were quantified in adipose tissue macrophages by flow cytometry. % of macrophages with F) TNF and G) IL-6 were quantified at baseline and following 8-hour activation with 50 ng/mL lipopolysaccharide (LPS) ex vivo. Means ± SEM, *p<0.5, **p<0.1, ***p<0.01, ****p<0.001 by one-way ANOVA and post-hoc analysis.

## 4. Discussion

Weight cycling worsens diabetes risk beyond that of stable weight obesity. Previous studies from our groups suggest that diet-induced weight loss drives immune phenotypes that may contribute to worsened glucose tolerance upon weight regain. This work investigated whether exercise during weight-loss could reduce exhausted CD8+ T cell accumulation in the adipose tissue and mitigate glucose intolerance following weight regain. Interestingly, we found that exercise could attenuate CD8+ T cell exhaustion in the adipose tissue. This effect was remarkably sustained even following the cessation of exercise and weight regain. However, exercise during weight loss did not improve glucose tolerance after weight regain. Further investigation found that exercise did not reduce macrophage memory or cytokine production in response to weight loss and weight regain.

We were particularly interested in examining the impact of exercise on CD8+ T cells and glucose tolerance with weight cycling, because it is well established that CD8+ T cells expand^29^, promote inflammation^9^, and become exhausted^10^ in adipose tissue during obesity. Our group has further shown that weight loss drives CD8+ T cell clonal expansion ^14^ and exhaustion^12^. Consistent with these findings, we observed that obesity increased total and memory CD8+ T cells along with exhaustion markers such as PD-1 and TOX, which were further increased with weight loss. In support of our hypothesis, the addition of exercise during weight loss reduced the expression of exhaustion markers in adipose tissue CD8+ T cells by sequencing. While there was variability in the percent and number of exhausted CD8+ T cells measured by flow cytometry, exercise volume (km/day) was significantly associated with a reduction in exhausted CD8+ T cells following weight loss. This data suggests that exercise volume may influence adipose CD8+ T cell phenotypes. To our knowledge, this is the first study demonstrating that exercise reduces exhausted CD8⁺ T-cell populations in adipose tissue during weight loss and is consistent with evidence that immune responses to exercise depend on exercise volume and intensity^30^.

Importantly, exercise during weight loss reduced total, memory, and exhausted CD8+ T cells in adipose tissue following weight regain, despite a 9-week period during which animals no longer had access to running wheels. The concurrent reduction in both memory and exhausted CD8+ T cells suggests that exercise may not simply reverse the exhausted phenotype of existing T cells but rather reduce the activation, expansion, and chronic stimulation of these populations within adipose tissue. While we do not know the antigens that induce CD8+ T cell memory in the adipose tissue with weight gain and loss, it is possible that these antigens come from lipolysis or adipose remodeling with weight loss. If such, exercise may enhance the coupling of lipolysis and lipid utilization in working skeletal muscle and/or uniquely remodel the adipose microenvironment. Additionally, it is plausible exercise may increase the clearance of T cells from the tissue. Prior studies hypothesize that exercise reduces exhausted CD8+ T cells via mobilization into the bloodstream and apoptosis^16^. Another possibility is that exercise affects lymphatic circulation ^31^ that could favor T cell migration out of the tissue. Such mechanisms could contribute to the removal of dysfunctional T cell populations from adipose tissue, resulting in a lasting reduction in exhausted CD8+ T cells even after exercise cessation.

While we found a sustained impact of exercise on CD8+ T cells, exercise did not affect systemic metabolism or adipose macrophages following weight regain. Consistent with previous studies, we found that weight cycling further worsened glucose tolerance beyond that of one bout of weight gain. These results support both experimental models that have shown that repeated weight gain and loss worsen metabolic dysfunction^24,32,33^ and meta-analyses of human research indicating associations between weight cycling and risk of glucose intolerance^34^. Exercise is well established as an effective intervention for improving glucose tolerance and reducing the risk for type 2 diabetes^18,35^. However, in our study, voluntary wheel running during weight loss did not improve glucose tolerance following weight regain, suggesting that consistent exercise is important for metabolic protection. This aligns with data in humans showing that discontinuation of an exercise intervention results in a rapid loss of insulin sensitivity^36^. Furthermore, our study shows that differences in basal glucose levels were most pronounced between the lean and the exercised group. While differences among the obese, weight cycled, and weight cycled plus exercise groups were modest, they may be indicative of underlying shifts in adipose tissue inflammatory processes.

Unexpectedly, we found that exercise did not reduce macrophage inflammation or the development of innate immune memory. Macrophages are another cell population which contribute to adipose inflammation and play a central role in the development of insulin resistance and glucose intolerance in obesity^37^. Like many other studies^38,39^, we saw that TNF and IL-6 were increased in obesity. Our data also support prior studies from our group and others that have shown macrophage inflammation persists following weight loss^26,27^ and that macrophage cytokine production increases following weight regain^24^. In contrast to the improvements observed in CD8+ T cell exhaustion with exercise, exercise had no impact on macrophage cytokine production following weight regain. Previously, we and others have found that weight loss specifically drives innate immune memory in adipose macrophages^24,40,41^. In this study, we found that exercise had no effect on the expression of metabolic genes (*Tnf, Hk3, Ldha, Sdhb*), epigenetic histone modifications (H3K4me3, H3K27ac), or effector function defined by LPS-stimulated TNF and IL-6. Altogether, these findings suggest a limit to exercise as a stand-alone therapeutic for macrophage inflammation and glucose intolerance with weight cycling, but a significant and sustained impact on T cell function.

Our study is not without limitations. While the use of mice allows for control of genetics, environment, exercise opportunity, and diet, the application of this work to humans requires more study. The contribution of macrophages to weight cycling-associated diabetes risk in mice and humans also requires further testing. Moreover, we used male mice as they have been shown to have robust weight change on high fat diet and robust changes in glucose tolerance. Female mice do get worsened glucose tolerance with weight cycling, and we do not anticipate that exercise would have had different impacts by sex, however, this should be experimentally tested for confirmation. The use of voluntary wheel running also introduced variability in exercise exposure among the mice, however, this methodology illuminated the importance of exercise volume for changes in exhausted CD8+ T cell phenotypes. Additionally, a major strength of voluntary exercise is that it diminishes the stress response associated with forced exercise (e.g. treadmill or swimming). This is important because catecholamines and stress hormones are known to influence immune function^42^. There also may be differences in immune and metabolic outcomes with different types or timing of exercise. Specifically, whether exercise during the weight regain phase or resistance exercise impact glucose tolerance or macrophage inflammation could be explored. Our objective, however, was to test whether exercise had long lasting effects on immune responses to weight regain. If mice were given access to running wheels during weight regain, they gain remarkably less weight than physically inactive controls. Last, immune profiling was only done in the epididymal adipose tissue and therefore may not fully capture systemic immune adaptations occurring in other tissues like the spleen. While reductions in CD8+ T cell exhaustion did not restore systemic glucose metabolism in our mice, it is plausible that exercise-induced improvements in CD8+ T cell function could have a more beneficial impact on disease progression in cancer^43^, infection^44^ or autoimmunity^45^.

## 5. Conclusion

In summary, we found that exercise reduces exhausted CD8+ T cells, and particularly memory subsets, in the adipose tissue. This effect was sustained 9 weeks following the cessation of exercise and following weight regain. Moreover, while the addition of exercise to diet-induced weight loss did not prevent glucose intolerance following weight regain, our results support a role for adipose macrophage innate immune memory in metabolic dysfunction with weight cycling. Future studies should examine if therapeutic strategies targeting innate immune pathways can more effectively restore metabolic health in weight cycling and should further examine the use of exercise to reduce exhaustion of tissue resident CD8+ T cell populations in other chronic disease conditions.

## Acknowledgements

The VANderbilt Technologies for Advanced GEnomics core laboratory (VANTAGE) was used for single cell RNA-sequencing. Vanderbilt Metabolic Mouse Phenotyping Center [VMMPC (NIH DK135073; www.vmmpc.org)].

## Author Contributions

OKO, ECL, MAC, AHH, NCW, and HLC all contributed to the conceptualization of this work. OKO, MK, EN, CH, MAC, NCW, and HLC all contributed to the data collection, and OKK, ECL, MAC, NCW, and HLC contributed to data analysis. OKO wrote the initial draft of this work and OKO, ECL, NCW, and HLC contributed to data interpretation. All authors provided comments or edits and approved this work for submission.

