## Supplemental Figures for "Voluntary exercise during weight loss attenuates adipose CD8+ T cell exhaustion that persists after weight regain"

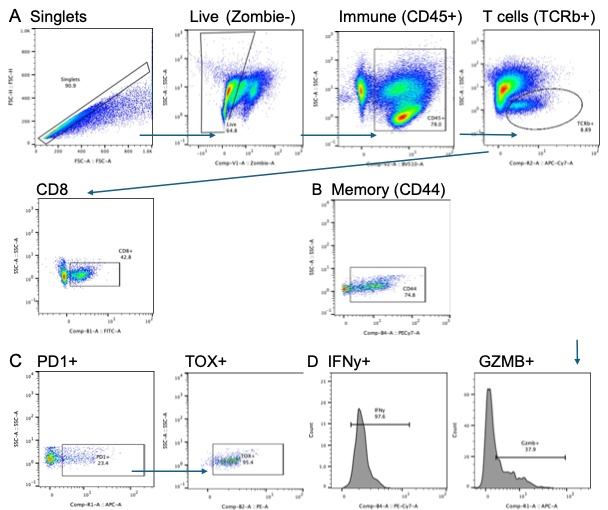

**Supplemental Figure 1: Adipose T cell gating.** A) CD8+ T cells were gated from singlets, live, CD45+, and TCRβ+ cells. B) CD44+ Memory cells were gated from CD8+ T cells. C) Exhausted T cells were gated as PD1+ and TOX+ from CD8+ T cells or Memory CD8+ T cells. D) IFNγ+ or GZMB+ were gated from CD8+ T cells.

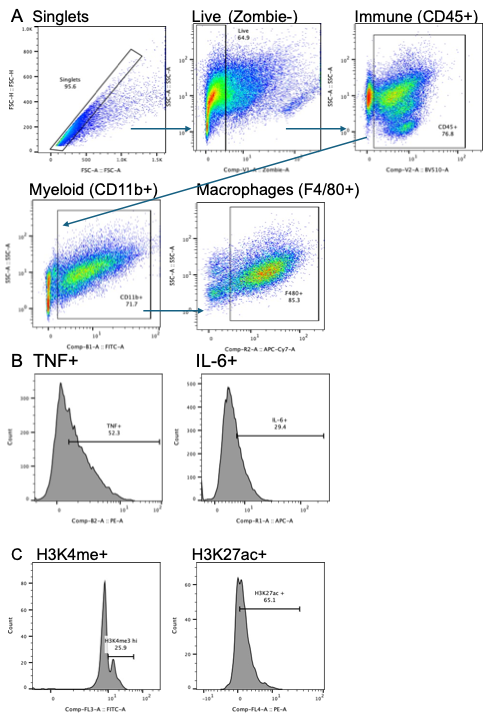

**Supplemental Figure 2: Adipose macrophage gating.** A) Macrophages were gated from singlets, live, CD45+, CD11b+, and F4/80+ cells. B) TNF+ and IL-6+ cells were gated from macrophages C) H3K4me3+ and H3K27ac+ cells were gated from macrophages.

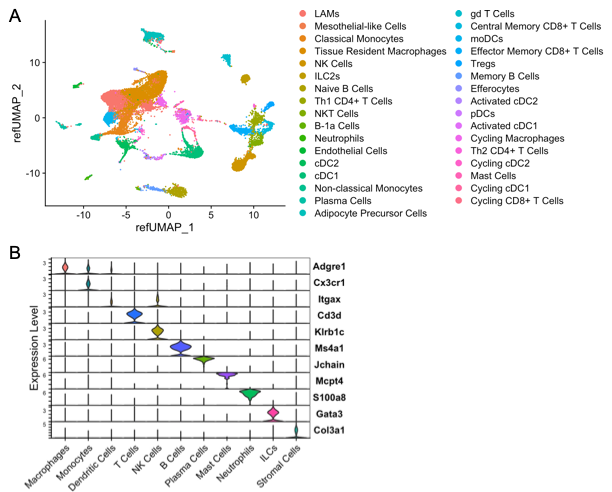

**Supplemental Figure 3: Adipose immune cell populations by single cell RNA- sequencing.** A) Unbiased clustering of 97188 single cells labeled broadly by cell type category and colored by high-resolution cell type identities via Uniform Manifold Approximation and Projection (UMAP). B) Clusters were named as in Cottam et al, 2022 and expression of selected genes support cluster names as displayed here.

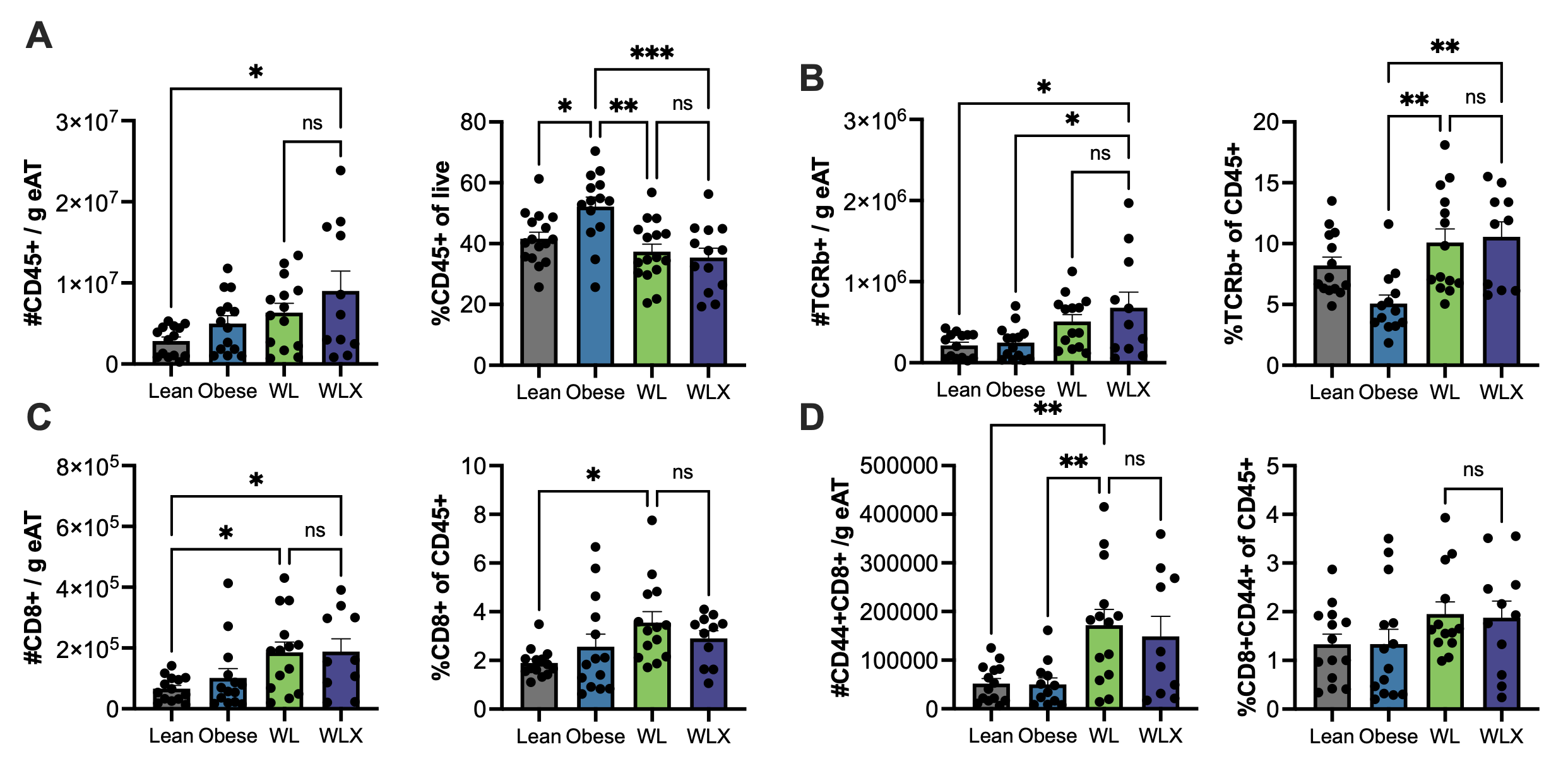

**Supplemental 4: All T cell populations by flow cytometry for our 18-week model.** # of cells/ gram of epididymal adipose tissue (AT) and % for A) CD45+ immune cells, B) T cells (CD45+TCRβ+), C) CD8+ T cells (CD45+TCRβ+CD8+), and D) CD8+ Memory T cells (CD45+, TCRβ+CD8+CD44+) for lean, obese, weight less (WL), and weight loss+ exercise (WLX) mice were quantified by flow cytometry. Means ± SEM, *p<0.5, ***p<0.01 by one-way ANOVA and post-hoc analysis.

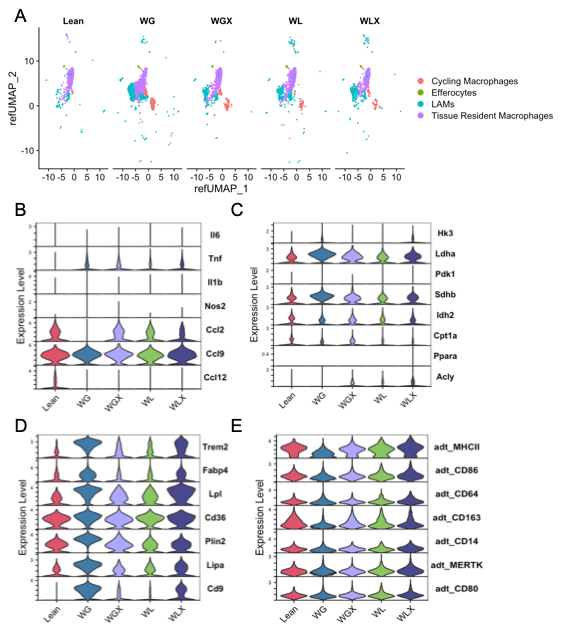

**Supplemental Figure 5: Exercise does not lessen the expression of inflammatory, glycolytic, or lipid handling genes with weight loss.** Macrophages populations were subset from the single cell RNA-sequencing data and A) a Uniform Manifold Approximation and Projection (UMAP) of these populations shows the distribution in lean, weight gain (WG), weight gain + exercise (WGX), weight loss (WL), and weight loss+ exercise (WLX) groups from Figure 1. Expression of B) inflammatory, C) metabolic, and D) lipid handling genes and E) inflammation related-protein surface markers are shown by group.
